# Full-season spring migration counts reveal seasonal contrasts in raptor migration through the eastern Black Sea flyway

**DOI:** 10.1101/2024.08.30.610513

**Authors:** Tohar Tal, Diego Jansen, Erik Jansen, Arthur W. Green, Elien Hoekstra, Marc Heetkamp, Filiep T’jollyn, Dries Engelen, Bart Hoekstra, Triin Kaasiku, Wouter M.G. Vansteelant

## Abstract

The extent to which geographical features like coastlines and mountain ranges funnel migrating birds depends on the seasonal context and direction of migration. Near Batumi, in the Republic of Georgia, the eastern Black Sea coast and Lesser Caucasus funnel over one million raptors through a 10-20 km wide coastal strip every autumn. The funnelling effect of the Lesser Caucasus appears much less evident for northbound migrants. Yet historical data suggest tens of thousands of raptors pass through the region in spring. To elucidate the composition and timing of spring raptor migration we conducted full-season migration surveys near Batumi in 2019, 2020 and 2022. In total, we recorded 33 species and, on average, counted 542,161 raptors (min. 455,799 - max. 618,848) annually. The bulk of the spring passage consisted of Black Kite *Milvus migrans* (239,649 ± 22,547) and Steppe Buzzard *Buteo buteo vulpinus* (194,029 ± 102,702). The most diverse and intense spring migration occurred from late March through mid-April, when the median passage date of 12 of 15 common species (>100 ind. y^-1^) occurred. Species with longer autumn migration periods tended to have longer spring migration periods, and most species had a longer migration period in spring than in autumn, likely due to larger age differences in timing during spring. Species’ abundance was up to an order of magnitude lower in spring than in autumn, consistent with a weaker bottleneck-effect in spring. Nevertheless, our results confirm the eastern Black Sea coast as a principal spring flyway and help redraw the map of East African-Eurasian migration for several Palearctic raptors. While we will not continue annual spring migration surveys, our data provides a baseline for detecting changes in raptor migration through short-term surveys and can help plan migration-based conservation and research at Batumi in spring.

## Introduction

Of all animal migrations, the mass migrations of raptors through so-called geographical bottlenecks are among the most accessible and visible events for the general public. These mass migrations arise from the funnelling effect of coastlines and mountain ranges that act as barriers or leading lines for diurnally migrating raptors seeking the safest, most energy-efficient routes toward their seasonal destinations (Kerlinger 1989, Bildstein 2006). Depending on the seasonal and wider geographical context, tens of thousands up to several million individuals of dozens of species may converge through a bottleneck within just a few weeks. Some of such bottlenecks in North America and Europe have been monitored for over half a century (Porter & Beaman 1985, Kerlinger 1989, Zalles & Bildstein 2000, Bildstein 2006), and new monitoring programs continue to be established at known and newly discovered bottlenecks around the world (Ruelas et al. 2000, Batista et al. 2004, Porras- Peñaranda et al. 2004, McCrary & Young 2008, Verhelst et al. 2011, Bayly et al. 2014, Fülöp et al. 2014, 2018, Heiss et al. 2020, Panuccio et al. 2021, Noby et al. 2022, Shi et al. 2023). Yet, there are still gaps in the migration monitoring network and much remains to be learned about raptors’ seasonal flyway use. Even within the East African-Eurasian flyway, used by several million raptors and where much historical research has been conducted (Porter & Beaman 1985, Shirihai et al. 2000), knowledge is still fragmentary in many places (Murgatroyd et al. 2021, Jobson et al. 2021). The seminal review of Shirihai et al. (2000) emphasised a particularly urgent need for spring surveys in the southeastern Black Sea region.

In general, more migrants are expected at bottleneck sites after the breeding season, when populations are inflated with large numbers of juvenile birds, of which many will perish before the following pre-breeding migration (Strandberg et al. 2009, Sergio et al. 2019, Serratosa et al. 2024, Mirski et al. 2024). However, the seasonal magnitude of migration at any given site also depends on the configuration of geographic features relative to the seasonal direction of migration. Bottlenecks with an hourglass configuration along a north-south axis such as the Central American Isthmus (Ruelas et al. 2000, Bildstein & Zalles 2001, Ruelas et al. 2009) or the Straits of Gibraltar (Evans & Lathbury 1973, Bensusan et al. 2007) exert a strong funnelling effect in both seasons. In other cases, the magnitude of migration may differ greatly between seasons at a given bottleneck depending on the defining geographic features’ orientation relative to the seasonal travel direction of birds, species-specific barrier-crossing abilities (Meyburg et al. 2017, Mellone 2020), seasonal time and energy constraints (Agostini et al. 2019), and other factors like seasonal weather conditions. For example, prevailing winds may exacerbate the barrier effect of a large body of water resulting in narrow-front migration in one season, while allowing a fast and direct water-crossing over a broad front in the other season (Panuccio et al. 2014, Nourani et al. 2016, Agostini et al. 2019). As a result, birds engage in seasonal loop migrations, preferring slightly different corridors within a given flyway, or even using entirely different flyways between seasons (Kerlinger 1989, Leshem & Yom-Tov 1996, Shirihai et al. 2000, Corso & Cardelli 2004, Katzner et al. 2016, Mirski et al. 2024). These and other factors like age-specific migration behaviours (Kjellen 1992, Schmid 2000, Sergio et al. 2014, Vansteelant et al. 2017, Phipps et al. 2019, Nourani et al. 2020) make that seasonal migration patterns can vary widely between sites (Kerlinger 1989, Bildstein 2006). Even today we lack basic information about the magnitude, timing, and composition of raptor migration in one or both seasons at key bottlenecks (Murgatroyd et al. 2021, Jobson et al. 2021).

In autumn, the Batumi bottleneck is one of few sites in the world where over one million raptors can be seen in a single season. The eastern Black Sea coast and the northern edge of the Lesser Caucasus make up a very pronounced geographical bottleneck for southbound migrants coming from the Kolkheti lowlands, which becomes progressively narrower from north to south until the mountains reach the sea just near the city of Batumi (Andrews et al. 1977, Gálvez et al. 2005, Verhelst et al. 2011, Fig. 1a-b). The extent to which raptors concentrate along the coast differs between species, but the use of the coastal route is intensified by dense cloud cover over the mountains limiting the development of thermal updrafts further inland (especially for obligate-soaring migrants in early autumn), and by easterly winds (especially for facultative-soaring species) (Vansteelant et al. 2014). The Batumi Raptor Count was established to monitor the autumn migration in 2008 (Verhelst et al. 2011), and a consistent survey effort since 2011 has revealed major changes in certain raptor populations using this flyway (Wehrmann et al. 2019, Vansteelant et al. 2020). Yet, the spring migration has only been roughly quantified in the past (Abuladze 2013).

**Fig. 1.**
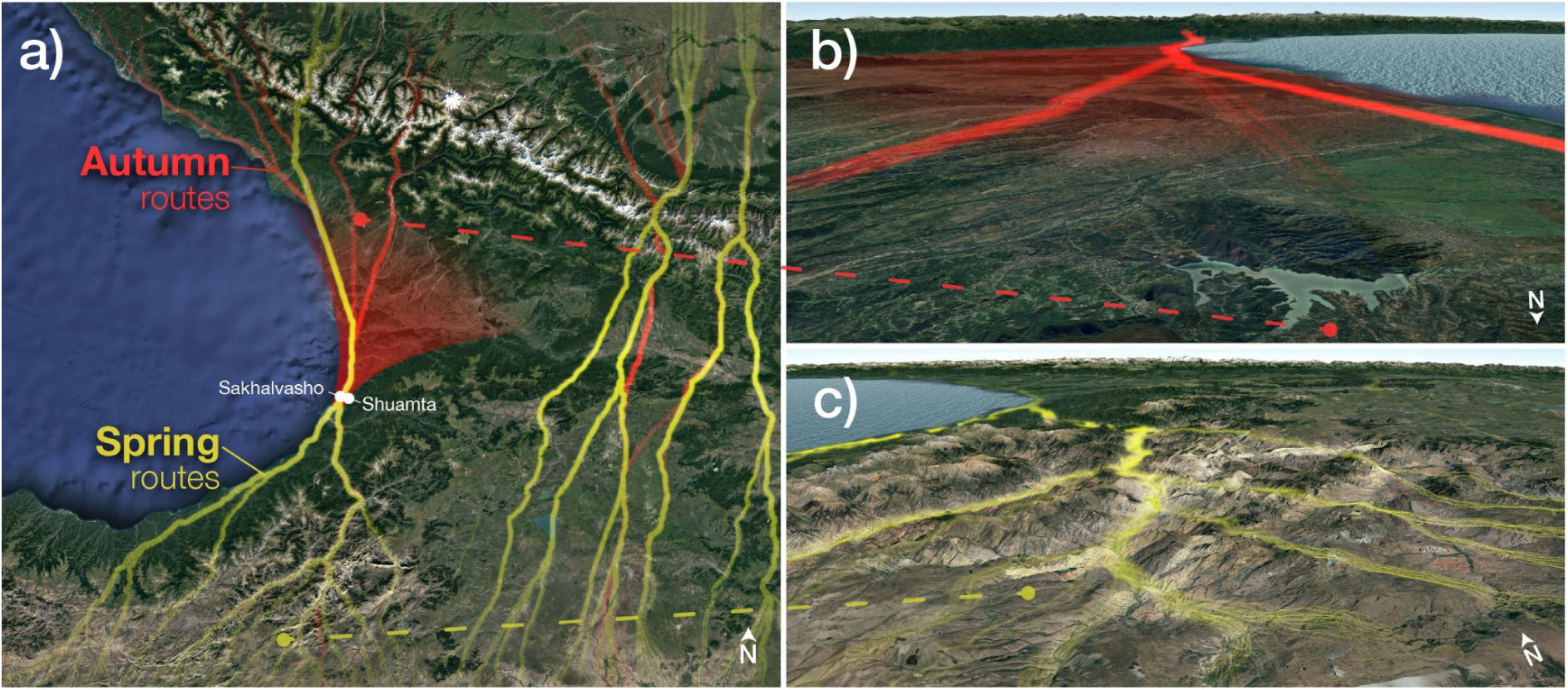
View of the Batumi bottleneck (a) from above, (b) looking south over the Kolkheti lowlands towards the bottleneck, and (c) north from the Armenian highlands towards the bottleneck. White dots indicate the Batumi Raptor Count watch sites of Sakhalvasho (2km from the coast at 324m asl) and Shuamta (6km from the coast at 414m asl) located within a 12km wide bottleneck. Only the Sakhalvasho site was used during the spring surveys. Routes are the shortest paths with the least elevation changes through the Caucasus region (a) and in particular the Batumi bottleneck (b and c). These routes illustrate the likely paths of convergence of birds if they made decisions only based on topography. Note how in autumn (b) the eastern Black Sea coast and the northern edge of the Lesser Caucasus form a wide topographical funnel zone (shaded in red), converging in a narrow ‘bottleneck’ for birds coming from the north. In spring (c), only the coast acts as an obvious leading line into the bottleneck, though birds might be guided by topography through the valleys of northeast Turkey. Satellite and topography extracted via Google Earth.

In this study, we report the first full-season spring migration surveys of northbound raptors in the eastern Black Sea flyway, in the Republic of Georgia, in the Caucasus region (Fig.1). Compared to autumn, the Batumi ‘bottleneck’ is much less clearly topographically defined in spring. Northbound migrants coming from the Armenian highlands in spring are instead expected to ‘filter through’ the many mountain passes of the Lesser Caucasus towards the Kolkheti lowlands (Fig. 1a, c).

Furthermore, climatic factors that may modulate the topographical barrier effect, e.g. through the development of thermal or orographic updrafts, are presumably different in spring than autumn. At a larger scale, certain species or populations that migrate through the eastern Black Sea flyway in autumn may use other flyways in spring, and vice versa (Shirihai et al. 2000, Pannucio et al. 2014, Vansteelant & Agostini 2021). Scant information about the spring migration of raptors across the Caucasus indicates that the spring migration through the eastern Black Sea flyway is much smaller in magnitude than the autumn migration, but presumably still of global relevance for several species (Shirihai et al. 2000, Abuladze & Edisherashvili 2003, Gálvez et al. 2005, Abuladze 2013).

After 15 years of autumn surveys by the Batumi Raptor Count (BRC, www.batumiraptorcount.org) and much speculation about what spring migration might look like, we decided to find out by conducting three pilot spring surveys at Batumi, Georgia, in 2019, 2020, and 2022. Our primary aims were to (1) describe the magnitude of migration and species composition, (2) quantify the timing and migration duration of “relatively common migrants” (i.e. average count > 100 ind.) and (3) compare the magnitude and duration of migration periods between the spring and autumn seasons within the Batumi bottleneck.

Overall, we expect the spring migration of raptors to be of substantially lower magnitude than in autumn due to (i) lower numbers of young birds in spring and (ii) a weaker funnelling effect of the Lesser Caucasus. However, based on BRC autumn surveys (Verhelst et al. 2011) and other spring surveys in the East African-Eurasian flyway (Shirihai et al. 2000), we expect a similar diversity of raptors in spring as in autumn, with 25-30 species occurring annually, and potentially still up to hundreds of thousands of Black Kite *Milvus migrans*, Steppe Buzzard *Buteo buteo vulpinus* and Honey Buzzards *Pernis apivorus*. Focusing on those species that are common and for which we achieve full coverage in both seasons, we expect most species to have a shorter core migration period in spring when adults are on their way to initiate breeding in a timely fashion, than in autumn when departure times can vary depending on individual breeding success, moult, and availability of local food resources (Kokko 1999, Newton 2023). At the same time we expect a longer total migration period due to a late and protracted migration of immature birds in spring (Leshem & Yom-Tov 1996, Yosef et al. 2003, Phipps et al. 2019, Mirski et al. 2024). We close by putting our observations in a larger context of the East African-Eurasian flyway and of historical data on the timing of spring raptor migration in Georgia, and considering the monitoring and conservation potential at Batumi in spring.

## Methods

### Study area, period, and species

The spring counts were conducted from a single count site located NE of Batumi along the west coast of Georgia, in the village of Sakhalvasho (41.68266°N, 41.72906°E). This site is situated on the same hill used during the BRC autumn surveys (Wehrmann et al. 2019) but on the south-facing side. The count site is located at roughly 325 m asl and 2 km east of the Black Sea coast in the foothills of the Lesser Caucasus mountains that rise to elevations >1200 m asl within 5-6 km from the coast (Fig.1). We conducted daily counts from 21 March until 31 May in 2019, and from 1 March until 26 May in 2020 and 2022. The difference in count period is a result of our observations in 2019, where upon arrival on 20 March there already seemed to be substantial raptor migration. By starting 20 days earlier we aimed to ensure full coverage of early-migrating species in 2020 and 2022. During the surveys, we counted all raptor species and those non-raptor species that are systematically recorded in autumn (Wehrmann et al. 2019).

### Data recording and processing

The count method used during the pilot spring surveys was a simplified version of our highly standardised autumn protocol (Wehrmann et al. 2019). The size of the team was significantly smaller with one to eight active counters on-site surveying from one (instead of two) count site(s). Counters were equipped with tally counters, binoculars, spotting scopes, and optionally a camera suitable for bird photography.

Spring counts were more easily interrupted by periods of inclement weather, such as heavy rain or snow, given the absence of any shelter on our pilot count site. This was for instance the case during the first three weeks of March 2022. Due to a constant snowfall resulting in snow cover >1 m deep we were often unable to reach the count site – although observations from our nearby guesthouse indicated migration activity was also minimal during that period. In 2019, we counted from a more inland location, but with a similar view as the main count site, on two days (41.68700°N, 41.77931°E). In 2020, due to COVID-19 quarantine measures set by the local government, we were forced to count from the garden of our guesthouse in the lower-lying village of Sakhalvasho (41.68657°N, 41.73352°E) for four consecutive days (26^th^ – 29^th^ March). The view from the terrace was sufficiently representative of our count site to conduct the daily counts, given that it was only 400 m northeast. During these days, except for the 27^th^ on which 32.000 birds (approximately 4% of the season total) were counted, the pace of migration was rather slow due to bad weather conditions.

As in autumn, we digitally recorded numbers of birds, with as much additional data as possible on species, age, sex, and colour morphs, using a tablet equipped with a modified version of the Trektellen app (Trektellen.org, Wehrmann et al. 2019). Higher-order taxonomic and morphological groups were used for birds that could not be identified to species level, such as the groups ‘medium-sized raptors’ and ‘large eagles’. In our autumn analyses a post-hoc correction is performed to estimate the proportions of species among unidentified birds on each day (Wehrmann et al. 2019, Vansteelant et al. 2020). However, due to higher numbers of unidentified raptors compared to identified raptors in our spring data, we were unable to develop an elegant correction of daily totals for our spring data. Therefore, to make seasonal comparisons, we also used the uncorrected daily species totals for autumn.

### Quantifying phenology and migration duration

We calculated seasonal totals and phenological metrics for all relatively common migrants. The seasonal timing was assessed based on quantile passage dates (5, 25, 50, 75, and 95%). We then calculated the duration of the main migration period (i.e. central 90%) as the period spanning from the Q5% to the Q95% quantile passage dates and of the core migration period (i.e. central 50%) as the period between the Q25% and the Q75% quantile passage date. As an additional metric, we calculated what proportion of individuals was seen on the biggest peak day of each season for each species. We calculated these metrics for spring as well as autumn to allow for seasonal comparisons.

In 2020 and 2022 – years in which the spring counts began on the 1st of March – we assume to have covered the full migration period of all recorded species as the migration only slowly started to pick up after the start of the count. However, this was not true for 2019, when the counts started on the 21st of March. This could lead to substantial bias in phenological metrics of the earliest spring migrants, especially the Q5%-passage date and the duration of the main migration period. To determine for which species the late 2019 start could be most problematic, we assessed what percentage of each species passed before March 21st in 2020 and 2022. If this exceeded 3%, we excluded 2019 from the calculation of phenological metrics of that species. A stricter percentage was found too exclusive for species with relatively low season totals of a few hundred birds. No corrections were needed for calculating the end of the migration periods, as migration of all species ceased before the end of the count in all years.

### Comparing spring and autumn abundance and phenology

We used the autumn data of 2018, 2019, and 2021 to compare seasonal abundance and migration periods for those species for which we achieved complete coverage in both seasons. Comparing spring totals to the immediately preceding autumn allows for cleaner comparisons because this accounts for the fact that some species changed significantly in abundance since the start of BRC’s autumn surveys (Vansteelant et al. 2020). We employed a two-sided unpaired t-test to determine whether seasonal differences in the magnitude and duration of main and core migration periods were statistically significant. Furthermore, we used a single-effect linear regression model to test whether the relative timing (Q50% passage date), the average duration of species’ main and core migration periods, and the proportion of birds seen on peak days were correlated between spring and autumn.

All statistical testing and data visualisation was conducted in R v.4.2.2 (R Core Development Team). The daily spring and autumn data and code to reproduce our analyses are publicly available through xxx and the BRC GitHub page.

## Results

We observed on average 542,161 raptors (min. 455,799 - max. 618,848) per spring across a total of 33 species (Table 1). Spring migration was dominated by Black Kite (239,649 ± 22,547), Steppe Buzzard (194,029 ± 102,702) and Honey Buzzard (54,994 ± 29,978), with an additional six species with average totals of 1,000 - 10,000 ind. and another six species with average totals 100 - 1,000 ind. For four of the 15 most common species (Imperial Eagle *Aquila heliaca*, Hen Harrier *Circus cyaneus*, Steppe Eagle *Aquila nipalensis* and Marsh Harrier *Circus aeruginosus*) we likely missed the first 3% of the spring migration in 2019 by starting 20 days later than in 2020 and 2022. For these species, we excluded 2019 from our calculations (Table 1). Of the 15 common species, 12 species had a median passage date between late March and mid-April, and the remaining three common species had median passage dates in late April and early May (Table 1, Fig. 2).

**Fig. 2.**
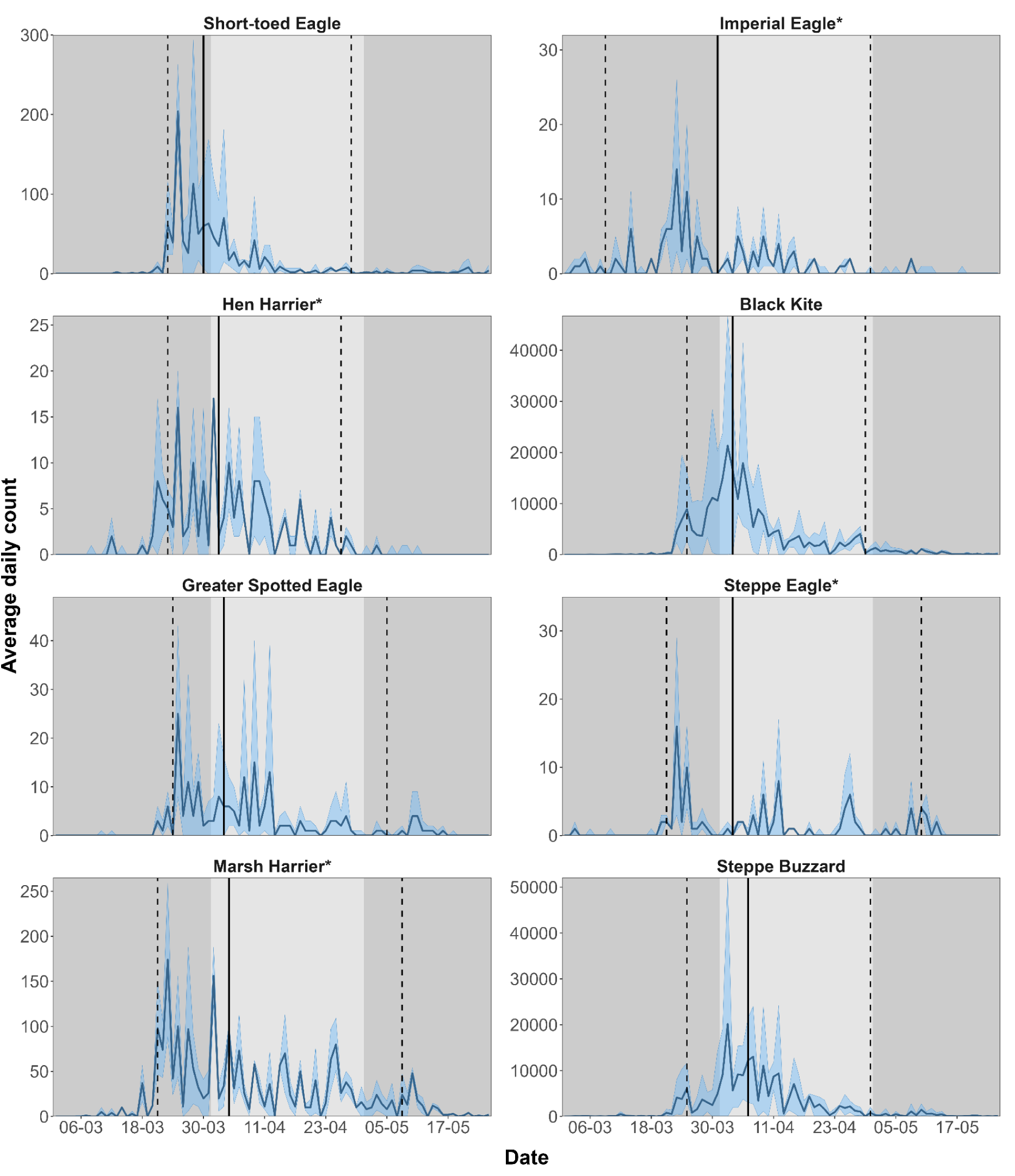
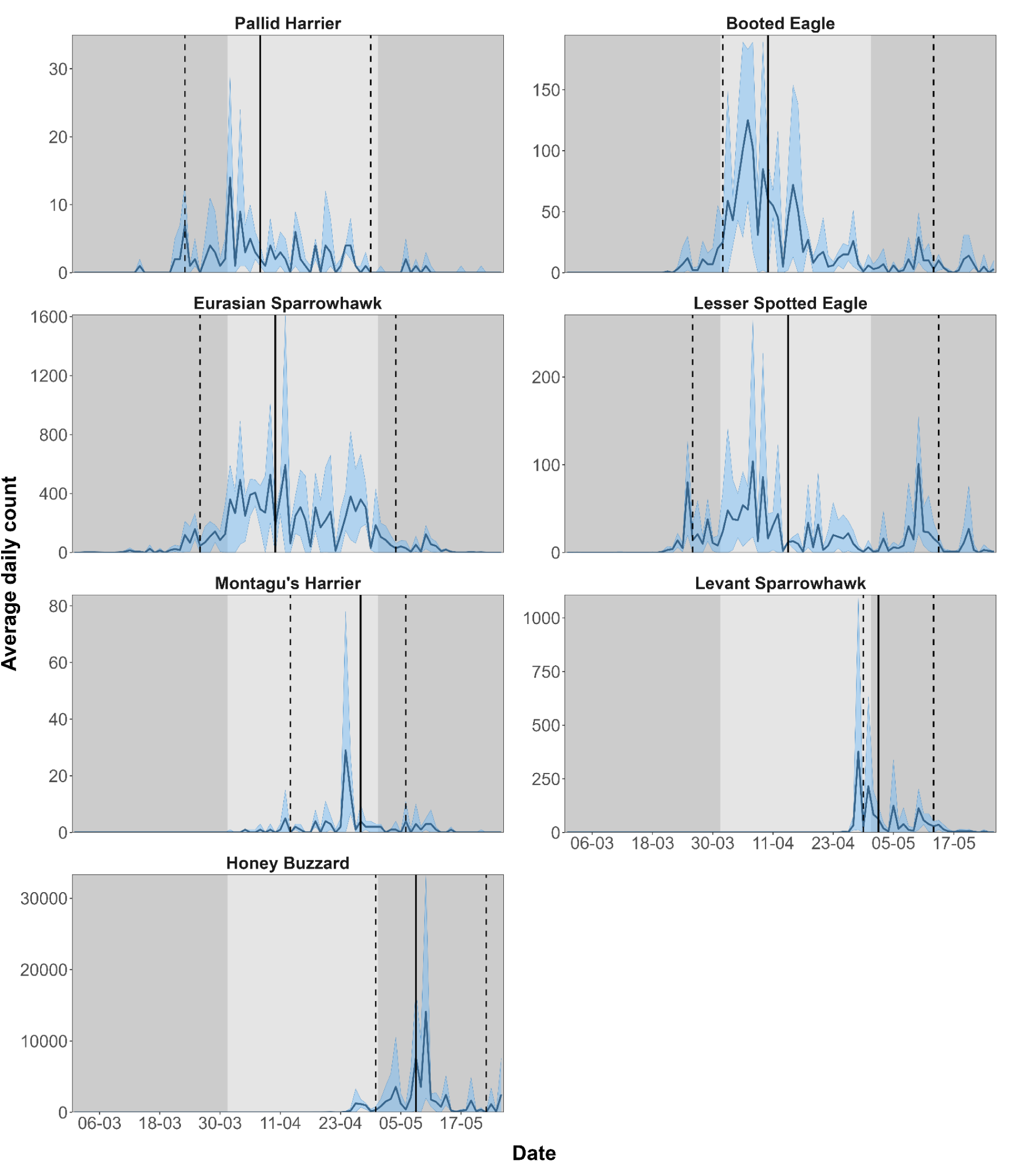
**a** Spring phenology of the first 8 of 15 most common raptors in spring (average total > 100 ind. yr-1). Dark blue lines show the mean daily count with the minimum and maximum count in light blue. Vertical lines indicate the median passage date (Q50%, solid) and the Q5% and Q95% quantile passage dates (dashed). Background shading indicates consecutive months: March (dark), April (light), and May (dark). * Early species: data based on 2020 and 2022 only. **b** Spring phenology of the last 7 of 15 most common raptors in spring (average total > 100 ind. yr-1). Dark blue lines show the mean daily count with the minimum and maximum count in light blue. Vertical lines indicate the median passage date (Q50%, solid) and the Q5%- and Q95% quantile passage dates (dashed). Background shading indicates consecutive months: March (dark), April (light), and May (dark).

**Table 1.**
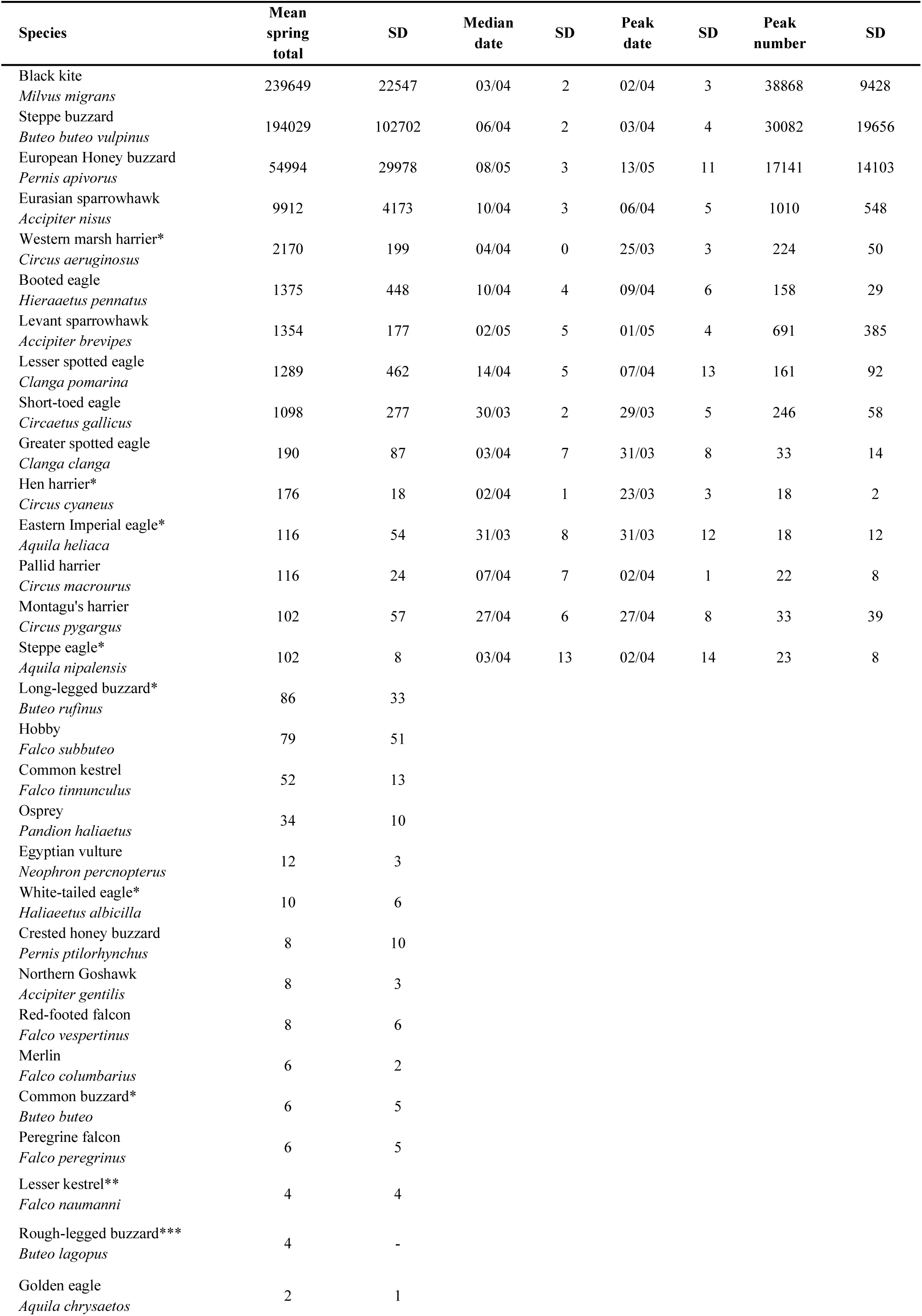
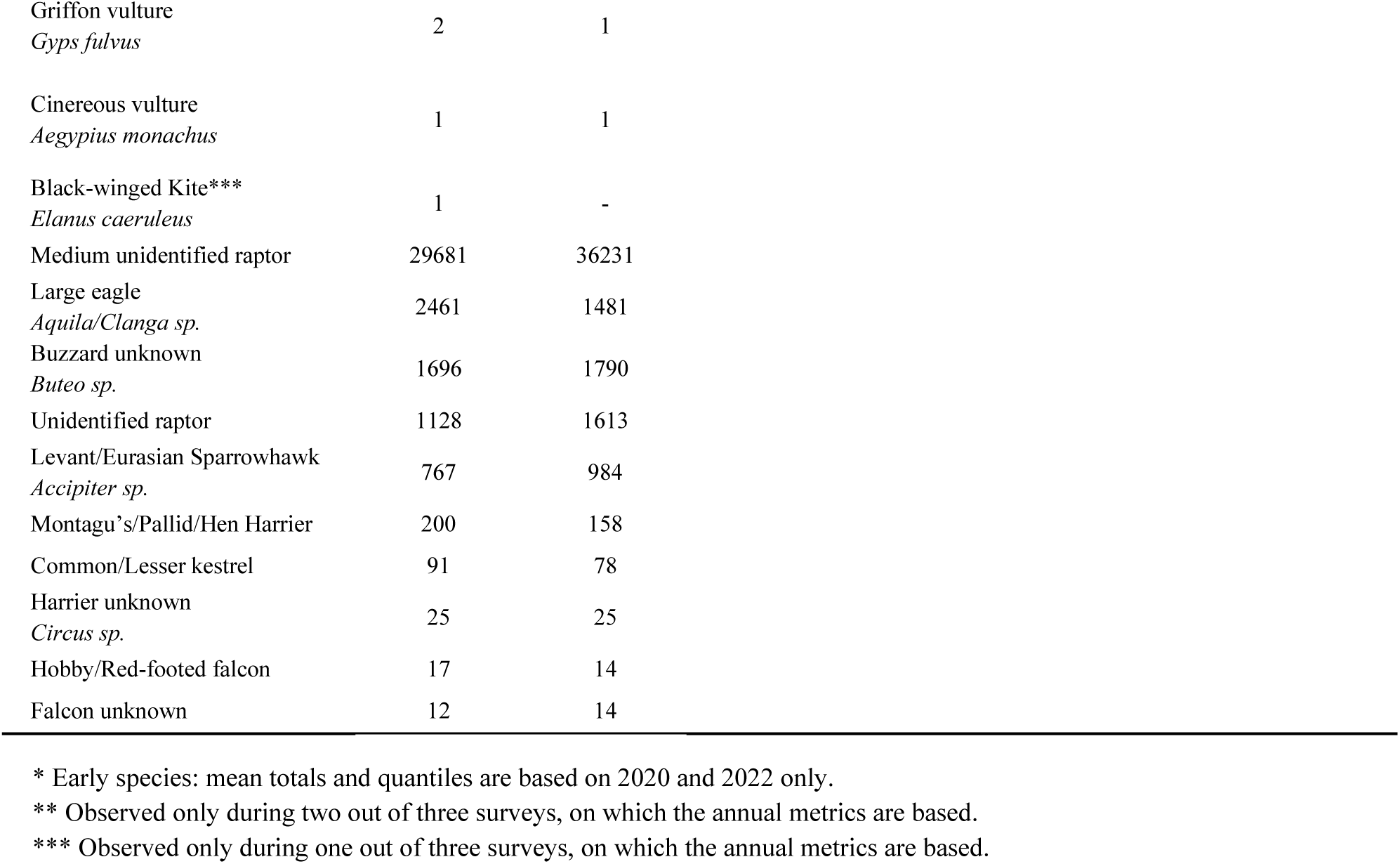
Overview of observed raptor species observed during the spring surveys in 2019, 2020 and 2022, including the mean annual total. For the relatively most common species (average total >100 ind.) we show the average median passage date, average date with the maximum day count, and the average number of individuals on the maximum day count.

| Species | Mean<br>spring<br>total | SD | Median<br>date | SD | Peak<br>date | SD | Peak<br>number | SD |
| --- | --- | --- | --- | --- | --- | --- | --- | --- |
| Black kite | 239649 | 22547 | 03/04 | 2 | 02/04 | 3 | 38868 | 9428 |
| *Milvus migrans* |  |  |  |  |  |  |  |  |
| Steppe buzzard | 194029 | 102702 | 06/04 | 2 | 03/04 | 4 | 30082 | 19656 |
| *Buteo buteo vulpinus* |  |  |  |  |  |  |  |  |
| European Honey buzzard | 54994 | 29978 | 08/05 | 3 | 13/05 | 11 | 17141 | 14103 |
| *Pernis apivorus* |  |  |  |  |  |  |  |  |
| Eurasian sparrowhawk | 9912 | 4173 | 10/04 | 3 | 06/04 | 5 | 1010 | 548 |
| *Accipiter nisus* |  |  |  |  |  |  |  |  |
| Western marsh harrier* | 2170 | 199 | 04/04 | 0 | 25/03 | 3 | 224 | 50 |
| *Circus aeruginosus* |  |  |  |  |  |  |  |  |
| Booted eagle | 1375 | 448 | 10/04 | 4 | 09/04 | 6 | 158 | 29 |
| *Hieraetus pennatus* |  |  |  |  |  |  |  |  |
| Levant sparrowhawk | 1354 | 177 | 02/05 | 5 | 01/05 | 4 | 691 | 385 |
| *Accipiter brevipes* |  |  |  |  |  |  |  |  |
| Lesser spotted eagle | 1289 | 462 | 14/04 | 5 | 07/04 | 13 | 161 | 92 |
| *Clanga pomarina* |  |  |  |  |  |  |  |  |
| Short-toed eagle | 1098 | 277 | 30/03 | 2 | 29/03 | 5 | 246 | 58 |
| *Circaetus gallicus* |  |  |  |  |  |  |  |  |
| Greater spotted eagle | 190 | 87 | 03/04 | 7 | 31/03 | 8 | 33 | 14 |
| *Clanga clanga* |  |  |  |  |  |  |  |  |
| Hen harrier* | 176 | 18 | 02/04 | 1 | 23/03 | 3 | 18 | 2 |
| *Circus cyaneus* |  |  |  |  |  |  |  |  |
| Eastern Imperial eagle* | 116 | 54 | 31/03 | 8 | 31/03 | 12 | 18 | 12 |
| *Aquila heliaca* |  |  |  |  |  |  |  |  |
| Pallid harrier | 116 | 24 | 07/04 | 7 | 02/04 | 1 | 22 | 8 |
| *Circus macrourus* |  |  |  |  |  |  |  |  |
| Montagu's harrier | 102 | 57 | 27/04 | 6 | 27/04 | 8 | 33 | 39 |
| *Circus pygargus* |  |  |  |  |  |  |  |  |
| Steppe eagle* | 102 | 8 | 03/04 | 13 | 02/04 | 14 | 23 | 8 |
| *Aquila nipalensis* |  |  |  |  |  |  |  |  |
| Long-legged buzzard* | 86 | 33 |  |  |  |  |  |  |
| *Buteo rufinus* |  |  |  |  |  |  |  |  |
| Hobby | 79 | 51 |  |  |  |  |  |  |
| *Falco subbuteo* |  |  |  |  |  |  |  |  |
| Common kestrel | 52 | 13 |  |  |  |  |  |  |
| *Falco tinnunculus* |  |  |  |  |  |  |  |  |
| Osprey | 34 | 10 |  |  |  |  |  |  |
| *Pandion haliaetus* |  |  |  |  |  |  |  |  |
| Egyptian vulture | 12 | 3 |  |  |  |  |  |  |
| *Neophron percnopterus* |  |  |  |  |  |  |  |  |
| White-tailed eagle* | 10 | 6 |  |  |  |  |  |  |
| *Haliaeetus albicilla* |  |  |  |  |  |  |  |  |
| Crested honey buzzard | 8 | 10 |  |  |  |  |  |  |
| *Pernis ptilorhynchus* |  |  |  |  |  |  |  |  |
| Northern Goshawk | 8 | 3 |  |  |  |  |  |  |
| *Accipiter gentilis* |  |  |  |  |  |  |  |  |
| Red-footed falcon | 8 | 6 |  |  |  |  |  |  |
| *Falco vespertinus* |  |  |  |  |  |  |  |  |
| Merlin | 6 | 2 |  |  |  |  |  |  |
| *Falco columbarius* |  |  |  |  |  |  |  |  |
| Common buzzard* | 6 | 5 |  |  |  |  |  |  |
| *Buteo buteo* |  |  |  |  |  |  |  |  |
| Peregrine falcon | 6 | 5 |  |  |  |  |  |  |
| *Falco peregrinus* |  |  |  |  |  |  |  |  |
| Lesser kestrel** | 4 | 4 |  |  |  |  |  |  |
| *Falco naumanni* |  |  |  |  |  |  |  |  |
| Rough-legged buzzard*** | 4 | - |  |  |  |  |  |  |
| *Buteo lagopus* |  |  |  |  |  |  |  |  |
| Golden eagle | 2 | 1 |  |  |  |  |  |  |
| *Aquila chrysaetos* |  |  |  |  |  |  |  |  |

|  |  |  |
| --- | --- | --- |
| Griffon vulture<br>*Gyps fulvus* | 2 | 1 |
| Cinereous vulture<br>*Aegypius monachus* | 1 | 1 |
| Black-winged Kite***<br>*Elanus caeruleus* | 1 | - |
| Medium unidentified raptor | 29681 | 36231 |
| Large eagle<br>*Aquila/Clanga sp.* | 2461 | 1481 |
| Buzzard unknown<br>*Buteo sp.* | 1696 | 1790 |
| Unidentified raptor | 1128 | 1613 |
| Levant/Eurasian Sparrowhawk<br>*Accipiter sp.* | 767 | 984 |
| Montagu's/Pallid/Hen Harrier | 200 | 158 |
| Common/Lesser kestrel | 91 | 78 |
| Harrier unknown<br>*Circus sp.* | 25 | 25 |
| Hobby/Red-footed falcon | 17 | 14 |
| Falcon unknown | 12 | 14 |

\* Early species: mean totals and quantiles are based on 2020 and 2022 only.
\*\* Observed only during two out of three surveys, on which the annual metrics are based.
\*\*\* Observed only during one out of three surveys, on which the annual metrics are based.

### Spring vs. autumn abundance

Of the three species that make up the bulk of migration in both seasons, the two early spring migrants - Steppe Buzzard and Black Kite - show no significant difference between spring and autumn in the magnitude of migration, with Black Kite spring counts being slightly, but consistently higher in each spring than in the preceding autumn (Table 2). All other common species (groups) except ‘medium-sized raptors’ and ‘large eagles’, occurred in significantly lower numbers in spring than in autumn (Table 2). In the most extreme cases, spring abundance was an order of magnitude lower than autumn abundance. Among common species, this was true for two of the three late spring migrants: Honey Buzzard (tens of thousands in spring vs. hundreds of thousands in autumn), and Montagu’s Harrier (100+ in spring vs thousands in autumn). Such a strong underrepresentation in spring was also observed for Pallid Harrier *Circus macrourus* and unidentified Montagu’s/Pallid/Hen harriers in general (at most a few hundred in spring versus thousands in autumn, Table 2).

**Table 2.** Comparison of the spring and autumn rounded mean annual total including results of the unpaired two-sided t-tests (<0.05 in *italics* and <0.01 in **bold**) for species of which we presume to have covered the entire migration period in both seasons.

| Species | <u>Mean annual count</u> |  |  |  | Significance |
| --- | --- | --- | --- | --- | --- |
|  | Spring | SD | Autumn | SD |  |
| Black kite<br>*Milvus migrans* | 239649 | 22547 | 217980 | 66727 | 0.639 |
| Steppe buzzard<br>*Buteo buteo vulpinus* | 194029 | 102702 | 331317 | 96672 | 0.167 |
| Honey buzzard<br>*Pernis apivorus* | 54994 | 29978 | 499263 | 14169 | **<0.001** |
| Western marsh harrier<br>*Circus aeruginosus* | 2170 | 199 | 8326 | 1687 | *0.022* |
| Booted eagle<br>*Hieraaetus pennatus* | 1375 | 448 | 6683 | 1348 | *0.014* |
| Lesser spotted eagle<br>*Clanga pomarina* | 1289 | 462 | 4085 | 421 | **0.002** |
| Short-toed eagle<br>*Circaetus gallicus* | 1098 | 277 | 1977 | 462 | *0.060* |
| Pallid harrier<br>*Circus macrourus* | 116 | 24 | 445 | 73 | *0.010* |
| Montagu's harrier<br>*Circus pygargus* | 102 | 57 | 1892 | 373 | *0.013* |
| Medium raptor<br>*Buteo/Pernis/Milvus* | 29681 | 36231 | 61490 | 29563 | 0.306 |
| Large eagle<br>*Aquila/Clanga* | 2461 | 1481 | 6582 | 2377 | 0.075 |
| Harrier unknown<br>*Circus sp.* | 200 | 158 | 4392 | 755 | **0.008** |
| Total – all species | 542162 | 81954 | 1153558 | 154570 | **0.008** |

We did not formally test seasonal differences in abundance for species (groups) that were uncommon (< 100 ind yr-1) in one or both seasons, or not systematically counted in recent autumn surveys.

However, a qualitative comparison reveals that all annual species for which we achieved full coverage during the comprehensive pilot autumn surveys 2008-2010, or during more recent autumn surveys, were less abundant in spring than autumn, in particular small falcons and Osprey *Pandion haliaetus* of which we recorded tens in spring vs. hundreds in autumn (Verhelst et al. 2011, Hoekstra et al. 2020). The few annual species that showed higher spring than autumn totals were all species for which we achieved much better spring than autumn coverage, such as Eurasian Sparrowhawk *Accipiter nisus*, Hen Harrier, Eastern Imperial Eagle and Long-legged Buzzard *Buteo rufinus* (Hoekstra et al. 2020).

Comparable spring and autumn maxima only occurred in non-annual species (e.g. Black-winged Kite *Elanus caeruleus* and Rough-legged Buzzard *Buteo lagopus*).

### Spring vs. autumn phenology

In general, the median passage data in spring was significantly and negatively correlated to the median passage data in autumn (β = -0.740, Std. error = 0.277, P-value = 0.032, R2 = 0.434). In other words: the latest spring migrants tended to be the earliest autumn migrants and vice versa (Fig. 3A). The percentage of birds passing on peak days (Fig. 3B) shows no clear correlation between spring and autumn (β = 0.273, Std. error = 0.486, P-value = 0.593, R2 = -0.094). Despite Fig. 3C - D suggesting clear positive correlations of main and core migration periods between spring and autumn, these were not significant (main migration period: β = 0.585, Std. error = 0.315, P-value = 0.105, R2 = 0.235; core migration period: β = 1.030, Std. error = 0.611, P-value = 0.136, R2 = 0.187).

**Fig. 3.**
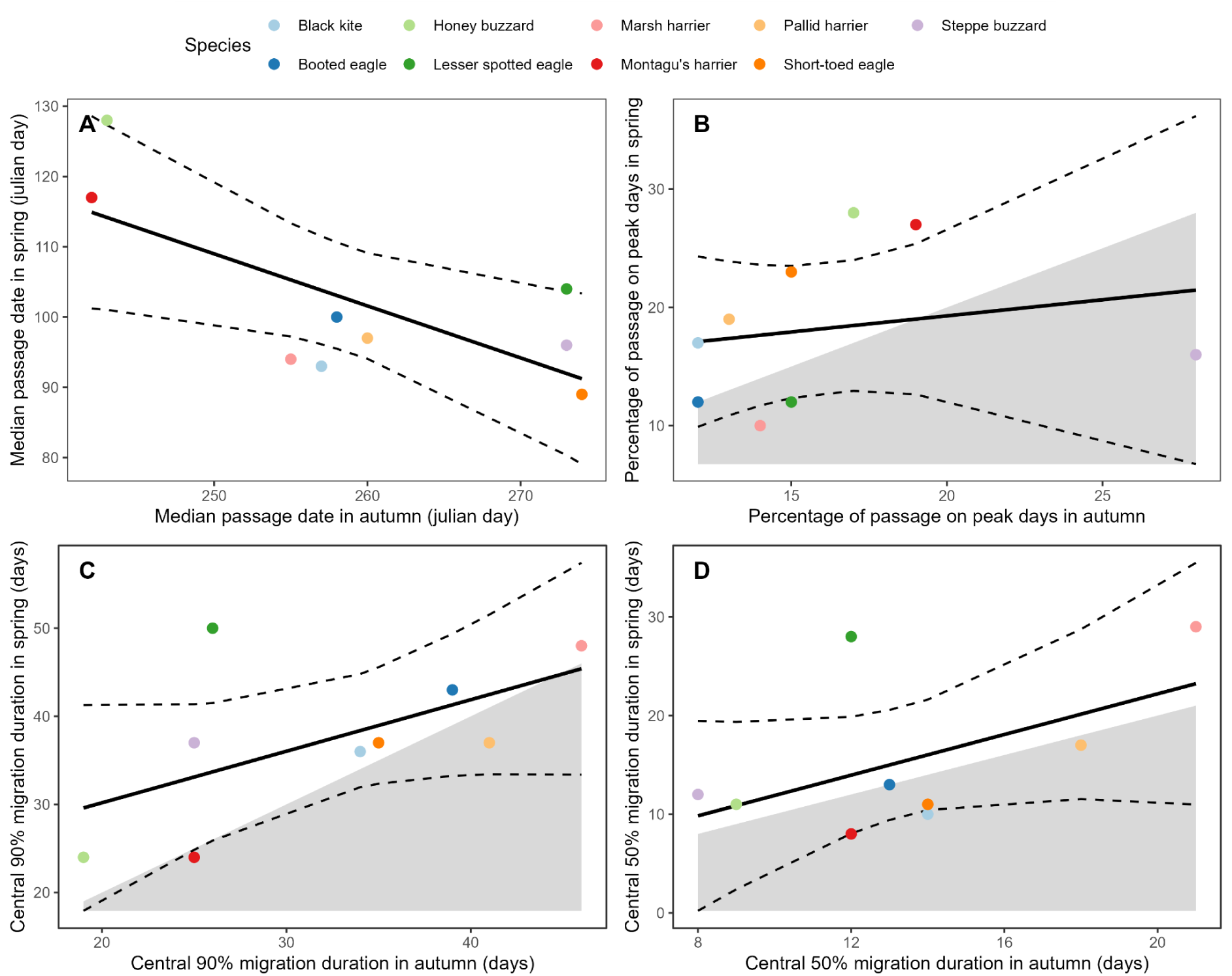
Seasonal comparison of the median passage date (A), percentage of birds passing on peak days (B), duration of the main 90% migration period (B), and duration of the core 50% migration period (D) for the 9 species with an average of >100 individuals per season and complete coverage during both spring and autumn. In all figures, a linear model is fitted (solid black line) including the lower and upper 0.95 confidence intervals (dashed black lines). In Figures A, B, and C we included the intercept of the x- and y-axis, and distinguished between autumn values higher than spring values (shaded) or autumn values lower than spring values (unshaded). Only the median passage date of species was significantly correlated between the spring and autumn seasons. For all other phenological metrics, we found no significant correlations.

The main migration period generally tended to last (slightly) longer in spring than in autumn (Fig. 3C, note 7 of 9 species in white area). While this difference was significant for only two species - Steppe Buzzard and Lesser Spotted Eagle *Clanga pomarina* (Table 3) - the relatively long main migration periods in spring seemed to be associated with very long tails of slow-paced migration in nearly all species (Fig. 2, note how the second half of the migration period from Q50% - Q95% generally spans much longer than the first half from Q5%-Q50%), much more than in autumn (note much less skewed phenological curves in Vansteelant et al. 2020). Furthermore, the only two species for which the main migration period was (not significantly) shorter in spring than autumn, Montagu’s Harrier and Pallid Harrier, would likely show longer spring main migration periods if we could account for the unidentified ringtail harriers that showed a much longer spring than autumn migration period (Table 3). Contrary to the main migration period, we found no general seasonal difference in the duration of species’ core migration periods or the percentage of birds passing on peak days (Fig. 3B, D; Tables 5- 6).

**Table 3.** Comparison of the spring and autumn rounded mean main migration duration (central 90%) in days including results of the unpaired two-sided t-tests (<0.05 in *italics* and <0.01 in **bold**) for species of which we presume to have covered the entire migration period in both seasons.

| Species | Length of central 90% (days) |  |  |  | Significance |
| --- | --- | --- | --- | --- | --- |
|  | Spring mean | SD | Autumn mean | SD |  |
| Black kite<br>*Milvus migrans* | 36 | 3 | 34 | 6 | 0.617 |
| Steppe buzzard<br>*Buteo buteo vulpinus* | 37 | 5 | 25 | 4 | *0.041* |
| Honey buzzard<br>*Pernis apivorus* | 24 | 8 | 19 | 3 | 0.4271 |
| Western marsh harrier<br>*Circus aeruginosus* | 48 | 4 | 46 | 4 | 0.484 |
| Booted eagle<br>*Hieraaetus pennatus* | 43 | 4 | 39 | 4 | 0.193 |
| Lesser spotted eagle<br>*Clanga pomarina* | 50 | 3 | 26 | 4 | **<0.001** |
| Short-toed eagle<br>*Circaetus gallicus* | 37 | 3 | 35 | 7 | 0.689 |
| Pallid harrier<br>*Circus macrourus* | 37 | 5 | 41 | 4 | 0.336 |
| Montagu's harrier<br>*Circus pygargus* | 24 | 6 | 25 | 2 | 0.658 |
| Medium raptor<br>*Buteo/Pernis/Milvus* | 41 | 2 | 28 | 9 | 0.115 |
| Large eagle<br>*Aquila/Clanga* | 48 | 1 | 22 | 2 | **<0.001** |
| Harrier unknown<br>*Circus sp.* | 42 | 3 | 28 | 2 | **0.002** |

### Other noteworthy observations

Every spring, we were struck by the strong tendency of Black Kites to aggregate along the coastline much more than in autumn, even making short-distance sea crossings (estimated 20 - 25 km) from roughly Batumi to Kobuleti, and frequently soaring several kilometres offshore. Additionally, each spring, a substantial portion of Black Kites passing the count site in mid-April would engage in aerial foraging (a behaviour that we never observed in autumn), opportunistically hunting small insects that swarmed between 11 - 18 April (Tal et al. 2021).

Throughout the spring seasons we recorded eight non-raptor species, summarised in Table S1. More anecdotal observations can be found in the spring migration reports published on the BRC website (Tal & Jansen 2019, Tal 2020, Hoekstra & Jansen 2022).

## Discussion

After establishing the significance of the Batumi bottleneck as one of the world’s major autumn raptor migration bottlenecks, and over a decade of standardised volunteer-based autumn surveys, we are now able to provide a detailed description of spring raptor migration through the eastern Black Sea Flyway. In line with our expectations, we recorded a similar diversity of raptors using the eastern Black Sea flyway in spring (33 species total) as in autumn (34 in 2008-2009 pilot surveys of Verhelst et al. 2011, and 38 species to date). As expected, 13 of the 15 common species for which we covered the full migration period in both seasons were much less abundant in spring than in autumn. Similar to our autumn surveys, the three most abundant species were Black Kites, Steppe Buzzards and Honey Buzzards, with the former two species showing a similar abundance with hundreds of thousands of individuals in both seasons. Contrastingly, the numbers of Honey Buzzards were an order of magnitude lower in spring (54,994 ± 29,978) than in autumn (499,263 ± 14,169). In general, the earliest spring migrants tended to be the latest autumn migrants, and vice versa. While we could not detect significant seasonal differences in phenological metrics based on three years of spring and autumn data, a great majority of species showed consistently longer main migration periods in spring, characterised by a relatively long tail of slow-paced migration.

Because we ran spring surveys with a relatively small team from only one count site, and spring counts were more often interrupted by bad weather than autumn counts, we likely underestimated true abundance more in spring than in autumn. However, the seasonal differences we found in species’ abundances (and migration periods) are too large to be explained solely by differences in coverage and thus reflect true seasonal differences in species’ population size and regional or large-scale flyway use that are discussed in more detail below. Another downside of working with small teams is that we had less time to spare for ageing and sexing, especially at times of intense migration. Recording demographic data also proved to be challenging because age-related features were less pronounced after the non-breeding period. Feathers of second calendar-year birds, for example, can be faded, worn, or already replaced, making it harder to distinguish them from older immatures and adults in spring (Forsman 2016). As a result, contrary to previous autumn analyses, we were unable to account for demographic factors like age in our spring analysis (Wehrmann et al. 2019, Vansteelant et al. 2020).

### Seasonal contrasts in raptor abundance

That most raptors would migrate past Batumi in lower numbers in spring than in autumn was expected due to the smaller numbers of young birds in the spring populations, and a weaker funnelling effect of the Lesser Caucasus on northbound migrants. Moreover, except for Honey Buzzard, all species underrepresented by an order of magnitude in spring compared to autumn are relatively slender-winged species such as ringtail harriers, Osprey, and small falcons. These species are well adapted to sustained powered flight and much less prone than broad-winged raptors to concentrate at bottleneck sites (Spaar 1997, Bildstein 2006), including Batumi in autumn (Verhelst et al. 2011).

Focusing on more typical bottleneck species, the only two species that were equally abundant in spring and autumn -despite presumably smaller spring flyway populations- were both early-spring migrants: Steppe Buzzard and Black Kite (Fig. 2A). And while the remaining early-spring species did show significantly lower spring than autumn abundance, they still occurred in a similar order of magnitude in both seasons. Contrastingly, the only specialist soaring migrant for which spring abundance was an order of magnitude lower than autumn abundance was Honey Buzzard, the latest spring migrant of all. Interestingly, this species stands out by the very low number of juveniles seen at Batumi in autumn (<10%, Vansteelant et al. 2020). As such, the seasonal contrast in the abundance of Honey Buzzards cannot be explained by the smaller (immature) population in spring. Instead, the exceptionally strong underrepresentation of this late-spring species could arise if the funnelling-effect of the Armenian Highlands and Lesser Caucasus towards the Black Sea coast is weakened over the course of the spring season.

Species passing in early spring face much harsher conditions by crossing the Armenian Highlands and Lesser Caucasus far inland (Gyllin 1974) than they do along the milder, coastal climate of the Black Sea-route past Batumi. Low temperatures and extensive snow cover can increase the energetic cost of crossing the Lesser Caucasus in early spring, for example through thermoregulation and weaker thermals necessitating active flight (Bildstein 2006, Bohrer et al. 2012), and limiting opportunities for refuelling (Niles et al. 1996, Bildstein 2006, Liminana et al. 2007, Trierweiler 2010). The fact that we observed hardly any visible migration during the late and severe winter spell in the southeastern Black Sea region in March 2022 certainly suggests snow and frost can block or divert migration (pers. comm.). Through mid and late spring, the interior of the Lesser Caucasus might become less hostile due to the greening up of the region. The contribution of geographic and seasonal climatic factors to the concentration of northbound raptors into the eastern Black Sea flyway could be disentangled by tracking early and late-spring migrants.

### Batumi spring migration in a flyway context

Spring migrants coming from Africa enter the Middle East via Egypt into Israel, or via Djibouti into Yemen by crossing the Bab-el-Mandab strait. From the Middle East they are presumed to follow one of three principal routes back to their respective breeding ranges (Shirihai et al. 2000): (1) northwestward over Türkiye and into eastern Europe via the Bosphorus Strait, (2) northward across the Armenian Highlands and the Caucasus in which the eastern Black Sea flyway is located, or (3) northeastward via the Caspian Sea into Central Asia. Our findings reveal that the magnitude of migration along the eastern Black Sea flyway matches the expectations of Shirihai et al. (2000) for some raptor species but exceeds previous estimations for others.

Species like Lesser Spotted Eagles that predominantly breed west of the Black Sea in Central and Eastern Europe migrate en masse via the Bosphorus in both seasons (Meyburg et al. 2000, 2004, Meyburg & Meyburg 2009, Üner et al. 2010, Meyburg 2021). A relatively small minority of 5,000- 10,000 birds take the eastern Black Sea route in autumn (Verhelst et al. 2011, Vansteelant et al. 2020), which likely originate from the easternmost populations in Ukraine, Belarus, westernmost Russia, and the Caucasus region itself (at least several dozen breeding pairs) (Keller et al. 2020). We now confirm a few thousand Lesser Spotted Eagles also take this route in spring (Shirihai et al. 2000); a seasonal contrast that can likely be accounted for by considerable juvenile mortality during the non-breeding season and a weaker topographic barrier effect in spring.

For species with substantial breeding populations north of the Caucasus, our data give new insights into seasonal flyway use. For example, we can confirm the expectation of Shirihai et al. (2000) that similar numbers of Steppe Buzzard use the eastern Black Sea flyway in spring as in autumn (Table 2, Verhelst et al. 2011) - albeit much higher numbers than were assumed to use this flyway. The seasonal passage at Batumi is also much greater than what has been recorded in other flyways around the Black and Caspian Sea in either season (Ullman & Ullman, 2010, Heiss 2013, Stanciu et al. 2017, Panuccio et al. 2018, Talebi E., pers. comm.), and accounts for the vast majority of Steppe Buzzards entering the Middle East via Israel in spring (circa 300,000 individuals; Shirihai et al. 2000, Yosef et al. 2002). Black Kite is a species for which different authors have formulated contradicting expectations about spring flyway use (Shirihai et al. 2000, Abuladze 2013, Panuccio et al. 2014).

While Shirihai et al. (2000) suggested the majority of the flyway population uses the eastern Black Sea flyway in both seasons, our results reveal that the concentration of Black Kites in the eastern Black Sea Flyway is even greater in spring than in autumn (Table 2), with spring counts being slightly higher than autumn counts despite the flyway population being smaller in spring. Tracking data of Black Kites, satellite-tagged at non-breeding grounds in Israel, supports the hypothesis that kites engage in a more narrow-front coastal migration over Batumi in spring than in autumn (Efrat R., pers. comm.). Even though the number of Black Kite in BRC autumn surveys doubled from <100,000 to nearly 200,000 in the period 2011-2018 (Vansteelant et al. 2020), and reached close to 300,000 in recent years (BRC, www.batumiraptorcount.org/data), hindcasting this trend to early 21st century suggests that the importance of the eastern Black Sea flyway for Black Kites in spring has been underestimated in the past (Panuccio et al 2014).

The major seasonal difference in Honey Buzzard abundance at Batumi (Table 2) at first glance supports the suggestion of Shirihai et al. (2000) that relatively more individuals migrate west over the Bosphorus in spring. However, spring migration surveys along the western routes through Türkiye yielded strikingly low Honey Buzzard counts, even considering incomplete coverage (<10,000 individuals, Üner et al. 2010, Arslangündogdu et al. 2018, Altundag & Karatas 2020). As it seems unlikely that Honey Buzzards breeding in northeastern Europe and western Russia migrate east along the Caspian Sea, we suggest that most of the hundreds of thousands of Honey Buzzards entering the East African-Eurasian flyway in spring (e.g. 360,184 ± 231,062 at Eilat, Israel, Shirihai et al. 2000) do cross the Caucasus, albeit over a much broader front than in autumn (e.g. <1,000 individuals at Besh Barmag, Azerbaijan in 2012, Heiss 2013). As stated above, being one of the latest spring migrants, Honey Buzzards likely encounter relatively hospitable conditions over the Lesser Caucasus (see also Gyllin 1974). Similarly, for other (earlier) species with substantial breeding populations north of the Caucasus, the lower spring than autumn abundance at Batumi can be explained without the use of alternative flyways. Instead, the general proclivity for broad-front migration can account for the particularly severe seasonal contrasts in the abundance of slender-winged species like Montagu’s Harrier. The comparatively moderate underrepresentation of soaring migrants in both early and late spring, such as Short-toed Eagle *Circaetus gallicus*, Booted Eagle *Hieraaetus pennatus*, and Levant Sparrowhawk *Accipiter brevipes* falls within expectations assuming a smaller spring flyway population in combination with a weaker funnelling effect of the Lesser Caucasus. Indeed, to our knowledge, there is little to no evidence that any of these species show a stronger spring than autumn concentration along the western Black Sea or Caspian flyways (Üner et al. 2010, Michev et al. 2011).

Besides trans-Caucasian migration occurring over a broader front, raptors breeding in central Asia, such as Steppe Eagles and Pallid Harriers, might generally be less inclined to detour through the (western) Caucasus in spring than in autumn. While Steppe Eagle counts at Batumi were significantly lower in spring (Table 1) compared to autumn (average 610, Zaytseva et al. 2022), counts on the eastern side of the Caucasus, at Besh Barmag (estimated 798 ± 0, Heiss 2016), are higher than our spring and autumn counts, and also higher than autumn counts at the same site (estimated 250, Heiss 2016). This likely reflects a tendency of Steppe Eagles to take a more easterly course over the Caucasus to return to their relatively eastern breeding areas in spring. Nevertheless, the passage across both ends of the Caucasus represents a small minority of the total Steppe Eagle population passing through the East African-Eurasian flyway (15,192 ± 1,553 entering through Israel, Weiss et al. 2019, several thousand wintering on the Arabian peninsula, e.g. Keijmel et al. 2020). Tracking data of Steppe Eagles wintering in the Middle East and breeding in Central Russia (Meyburg et al. 2012, Javed et al. 2014, Karyakin et al. 2019, 2023), corroborates that the vast majority of Steppe Eagles breeding in Central Asia enter and leave the Middle East southeast of the Caspian Sea (Shirihai et al. 2000), where systematic autumn surveys have recently been established (Panuccio et al. 2018, Talebi E., pers. comm.). The few Central Asian birds that do cross the Caucasus in autumn are even fewer in spring. Tracking data of Pallid Harriers from Kazakhstan similarly showed that birds migrating across the Caucasus in autumn, returned to Central Asia via more easterly routes south of the Caspian Sea in spring (Terraube et al. 2012).

### Seasonal contrasts in duration of migration

Based on nine species for which we have data of comparable quality in both seasons, the main spring migration period lasted 24-50 days (Table 3) and the core migration period was 8-29 days (Table 4). Though the duration of species’ migration periods was not significantly correlated between seasons, species with longer core and main migration periods in autumn tended to have longer migration periods in spring (Fig. 3C-D). Such between-species differences in the duration of migration tend to be associated with life-history and functional traits such as migration distance, breeding area size, longevity, wing morphology (and associated flight mode), and the extent to which demographic groups differ in timing within species (Leshem & Yom-Tov 1996, Bildstein 2006).

**Table 4.** Comparison of the spring and autumn rounded mean core migration duration (central 50%) in days including results of the unpaired two-sided t-tests (<0.05 in *italics* and <0.01 in **bold**) for species of which we presume to have covered the entire migration period in both seasons.

| Species | Length of central 50% (days) |  |  |  | Significance |
| --- | --- | --- | --- | --- | --- |
|  | Spring mean | SD | Autumn mean | SD |  |
| Black kite<br>*Milvus migrans* | 10 | 3 | 14 | 5 | 0.293 |
| Steppe buzzard<br>*Buteo buteo vulpinus* | 12 | 4 | 8 | 5 | 0.352 |
| Honey buzzard<br>*Pernis apivorus* | 11 | 10 | 9 | 2 | 0.684 |
| Western marsh harrier<br>*Circus aeruginosus* | 29 | 5 | 21 | 6 | 0.179 |
| Booted eagle<br>*Hieraaetus pennatus* | 13 | 5 | 13 | 2 | 0.923 |
| Lesser spotted eagle<br>*Clanga pomarina* | 28 | 14 | 12 | 2 | 0.186 |
| Short-toed eagle<br>*Circaetus gallicus* | 11 | 6 | 14 | 1 | 0.364 |
| Pallid harrier<br>*Circus macrourus* | 17 | 3 | 18 | 9 | 0.910 |
| Montagu's harrier<br>*Circus pygargus* | 8 | 6 | 12 | 1 | 0.369 |
| Medium raptor<br>*Buteo/Pernis/Milvus* | 25 | 4 | 12 | 6 | *0.036* |
| Large eagle<br>*Aquila/Clanga* | 15 | 6 | 10 | 3 | 0.271 |
| Harrier unknown<br>*Circus sp.* | 23 | 5 | 13 | 1 | 0.068 |

**Table 5.** Comparison of the spring and autumn rounded mean percentage of passage on peak days including results of the unpaired two-sided t-tests (<0.05 in *italics* and <0.01 in **bold**) for species of which we presume to have covered the entire migration period in both seasons.

| Species | Percentage of passage on peak days |  |  |  | Significance |
| --- | --- | --- | --- | --- | --- |
|  | Spring mean | SD | Autumn mean | SD |  |
| Black kite<br>*Milvus migrans* | 17 | 5 | 12 | 4 | 0.333 |
| Steppe buzzard<br>*Buteo buteo vulpinus* | 16 | 6 | 28 | 6 | 0.071 |
| Honey buzzard<br>*Pernis apivorus* | 28 | 8 | 17 | 6 | 0.122 |
| Western marsh harrier<br>*Circus aeruginosus* | 10 | 3 | 14 | 6 | 0.496 |
| Booted eagle<br>*Hieraaetus pennatus* | 12 | 3 | 12 | 4 | 0.965 |
| Lesser spotted eagle<br>*Clanga pomarina* | 12 | 4 | 15 | 4 | 0.346 |
| Short-toed eagle<br>*Circaetus gallicus* | 23 | 3 | 15 | 3 | *0.039* |
| Pallid harrier<br>*Circus macrourus* | 19 | 6 | 13 | 5 | 0.246 |
| Montagu's harrier<br>*Circus pygargus* | 27 | 18 | 19 | 6 | 0.523 |
| Medium raptor<br>*Buteo/Pernis/Milvus* | 32 | 21 | 26 | 6 | 0.641 |
| Large eagle<br>*Aquila/Clanga* | 14 | 3 | 29 | 3 | **0.005** |
| Harrier unknown<br>*Circus sp.* | 14 | 4 | 23 | 3 | *0.040* |

Generally, adult birds are expected to exhibit a higher pace of migration, and potentially a more synchronised passage in spring compared to autumn in an attempt to reach the breeding grounds earlier and occupy high-quality territories (Kokko 1999, Nilsson et al. 2013). That the majority of common species showed (significantly) longer main migration periods in spring than autumn at Batumi seemingly suggests otherwise. However, as no difference is made between age classes in this study, this effect is likely due to a larger spread in the timing of different age groups during spring migration (Phipps et al. 2019, Weiss et al. 2019). Long tails typically accounted for a much larger portion of the main migration period than the early or core migration periods. Contrary to the main migration period (i.e. central 90% period) the duration of the core migration period (i.e. central 50% period) did not differ significantly between seasons in any species, which points more towards these core periods mostly involving adult breeders.

The seasonal difference in duration of the main migration period is by far most pronounced for the Lesser Spotted Eagle: a long-lived species with a protracted immature phase, in which juveniles and adults show strong overlap in autumn migration timing (Meyburg et al. 2017, Vansteelant et al. 2020), whereas immatures migrate much later than adults in spring (Meyburg et al. 2001). A similar seasonal pattern would be expected and also seems to emerge from our seasonal data, for other large and long-lived *Aquila* and *Clanga* eagles at Batumi. Unfortunately, we achieved insufficient coverage of these species in autumn to make a formal seasonal comparison. The seasonal difference in migration duration seems to be only small for species like Short-toed Eagle, Booted Eagle and especially Honey Buzzard (Mirski et al. 2024). These species show long tails in spring migration and tend to show delayed immature migration in spring as *Aquila* and *Clanga* eagles do, but also the autumn migration of juveniles is delayed by several days-weeks compared to adults (Vansteelant et al. 2017, Vansteelant et al. 2020). In other cases with relatively small seasonal differences in migration duration, like Black Kite, there is hardly any difference in the autumn timing of juveniles and adults - as for large eagles (Vansteelant et al. 2020) - but differently than large eagles also only limited age differences in timing during spring (Sergio et al. 2014).

### Current spring phenology in the context of historical data

Comparing the timing of various species in our spring surveys to historical spring migration data from the 1970s and 80s summarised by Abuladze (2013), it appears that the spring migration period of Black Kite has remained identical, with the main period occurring from late March to mid-April, with the peak in the first week of April. By contrast, Short-toed Eagles showed a substantially earlier spring migration in our surveys (peak at the end of March) compared to historical data (peak 10-20 April). As such, Short-toed Eagle is the only species for which our data suggest a substantial advance in timing may have taken place compared to several decades ago, which would be consistent with general expectations about how migrants ought to respond to climate change (Newton 2023).

Furthermore, short-distance migrants are expected to respond more quickly to climate change than long-distance, trans-Saharan migrants (Newton 2023), and (a seemingly growing) part of the Short-toed Eagle population spends the non-breeding season around the Mediterranean (Vansteelant et al. 2020). However, for almost all other species for which historical data are provided by Abuladze (2013), our data suggest that spring migration now occurs later in the season than it did several decades ago, regardless of the species’ non-breeding distributions. For example, Abuladze (2013) suggests that Steppe Buzzard already showed relatively strong movement during early March whereas we only saw substantial passage after March 20 with peak days in early April. Other examples, where our data indicate a substantially later peak and longer tails in spring migration periods, are Marsh *Circus aeruginosus*, Montagu’s and Pallid Harrier, Levant and Eurasian Sparrowhawk, Lesser Spotted Eagle and Booted Eagle. While these comparisons are fraught with numerous issues, the implied phenological changes would be quite large in some cases (up to 2-3 weeks) and cannot easily be dismissed, even if they run counter to general expectations.

### Implications for monitoring, conservation, and avitourism

The possibility to monitor juvenile numbers, and thus reproductive success, is often what makes post-breeding autumn migration the primary focus of long-term raptor migration surveys (Jobson et al. 2021). Even though our pilot surveys proved the eastern Black Sea coast to be a principal spring migration flyway for many Palearctic raptors, the potential for systematic monitoring of spring raptor abundance and demography seems more limited than in autumn. Nevertheless, the data presented here offers a useful baseline to detect long-term changes in raptor migration by repeating small-scale spring surveys every 6-10 years. Even though this would limit the power to detect only moderate to large changes (Lewis & Gould 2000), intermittent migration counts may still be used for population and conservation assessments of selected species (Weiss et al. 2019).

Furthermore, our pilot surveys provide great potential to expand conservation and environmental education efforts in the Batumi region during spring migration. In particular, the knowledge we gained about the intensity and timing of spring migration can help designing and planning migration-based education projects. This is relevant because schools in the region are mostly operating field excursions in spring, whereas schools are closed until far into the autumn migration.

Similar to autumn, the Batumi region has the potential to attract significant migration-based avitourism activity in spring. Aside from the spectacular migration of large eagles and Black Kites early in the season, our field experience suggests that the presence of non-raptor migrants throughout the season, both on the count site and in surrounding natural areas, is much more pronounced in spring than in autumn. Additionally, illegal bird hunting – which is a persistent issue that affects migrating raptors (Sándor et al. 2017, Sándor & Anthony 2018) and we know keeps many potential ecotourists from visiting Batumi in autumn – rarely takes place in spring (van Maanen et al. 2001, Sándor et al. 2017, pers. comm.). And while the absence of a dedicated monitoring team may make the site somewhat less attractive in spring, some potential visitors may prefer to visit outside the main avitourism season. The general housing availability remains similar to autumn and the current lack of infrastructure tailored to spring conditions (i.e. appropriately placed, sheltered observatory) could be remediated relatively easily by targeted investments.

## Conclusions

In conclusion, our full-season spring surveys yielded novel insights into the seasonal importance of the eastern Black Sea flyway for Palearctic raptor species. In fact, our publicly visible data has already been used to this end by several authors (e.g. McGrady et al. 2021, Meyburg 2021, Onrubia & Martin 2021, Stark & Liechti 2021, Väli & Mirski 2021). BRC does not deem it favourable to invest in a structural, annual spring monitoring. However, intermittent spring migration surveys may allow the detection of demographic and phenological changes in Palearctic raptor species. As migration survey and monitoring efforts continue to grow at known bottlenecks (Murgatroyd et al. 2021, Noby et al. 2022) and newly discovered sites (Panuccio et al. 2018, Jobson et al. 2021), and with new tracking studies underway, we can look forward to exciting discoveries on East African-Eurasian raptor migration in the near future.

## Acknowledgements

We are grateful for all the volunteers who have helped us on-site performing the spring counts, particularly: Aslan Bolkvadze, Bjorn Alards, Katharine Khamhaengwong, Theo Askov, Hans Henrik Schou, Leo Akhmeteli, Ron Lawie, Rosena Tomova, Marijn Prins, Leslie Freeman, Leo Ballering, Mohammad A. Tabari, Geoff Oliver, Nina Askov, Mehmet Deli. The BRC spring counts would also not have been possible without the support of Ruslan Lomadze and Rusiko Meladze, their families, and neighbours in the lovely village of Sakhalvasho. We thank Gerard Troost for revolutionising our data recording strategy by developing the database Trektellen.org with its migration count app.

Finally, we thank the Ornithological Society of the Middle East, the Caucasus and Central Asia (OSME) as the main funder of the spring counts.

## Supplemental Material

**Fig. S1.**
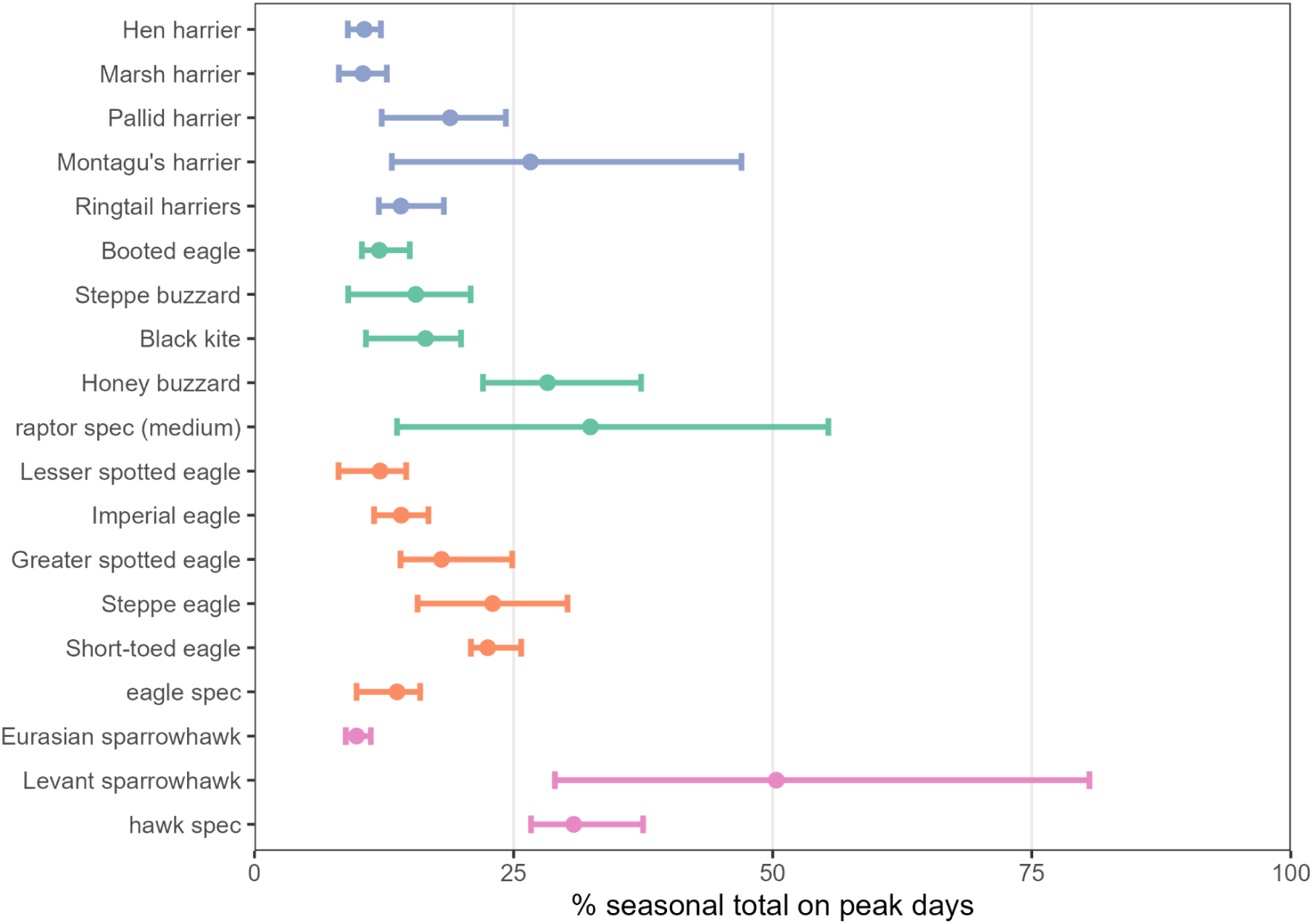
The mean percentage of birds seen on the biggest peak day of each season (dot). The whiskers indicate the minimum and maximum percentage recorded for each species in any of the years. Colors denote morphological groups (purple = harriers, green = medium raptors, orange = large eagles, and pink = sparrowhawks).

**Table S1.** Comparison of the spring and autumn rounded mean annual counts for non-raptor species. Including the standard deviation and the number of years (i.e. seasonal surveys) in which the species has been observed and on which the annual metrics are based.

| Species | Spring |  |  | Autumn |  |  |
| --- | --- | --- | --- | --- | --- | --- |
|  | Mean | SD | Years | Mean | SD | Years |
| Black stork<br>*Ciconia nigra* | 435 | 153 | 3 | 1732 | 181 | 3 |
| Eurasian crane<br>*Grus grus* | 282 | 186 | 3 | 248 | 73 | 3 |
| European roller<br>*Coracias garrulus* | 12 | 6 | 3 | 1534 | 530 | 3 |
| Stock dove<br>*Columba oenas* | 100 | 120 | 3 | 1188 | 607 | 3 |
| European turtle dove<br>*Streptopelia turtur* | 6 | 4 | 2 | 1354 | 1192 | 3 |
| Great white pelican<br>*Pelecanus onocrotalus* | 3 | - | 1 | 90 | - | 1 |
| White stork<br>*Circaetus gallicus* | 85 | 47 | 3 | 470 | 151 | 3 |
| Common wood pigeon<br>*Columba palumbus* | 433 | 197 | 3 | 752 | 756 | 3 |

